# Positive Feedback Regulation of *fzd7* Expression Robustly Shapes Wnt Signaling Range in Early Heart Development

**DOI:** 10.1101/2021.09.12.458649

**Authors:** Takayoshi Yamamoto, Yuta Kambayashi, Boni Afouda, Yuta Otsuka, Claudiu Giuraniuc, Tatsuo Michiue, Stefan Hoppler

**Author notes:** **Corresponding author:** Takayoshi Yamamoto, PhD. **Senior Authors:** Prof. Tatsuo Michiue.

## Abstract

Secreted molecules called morphogens govern tissue patterning in a concentration-dependent manner. However, it is still unclear how reproducible patterning can be achieved with diffusing molecules, especially when patterning differentiation of a thin region. Wnt is a morphogen that organizes cardiac development; especially Wnt6 patterns the cardiogenic mesoderm to induce differentiation of a thin pericardium in *Xenopus*. It is, however, unclear how Wnt6 can pattern such a thin tissue. In this study, we reveal that a Wnt receptor, *frizzled7*, is expressed in a Wnt-dependent manner in the prospective heart region, and that this receptor-feedback is essential for shaping a steep gradient of Wnt. In addition, the feedback imparts robustness against fluctuations of Wnt ligand production and allows the system to reach a steady state quickly. We also found a Wnt antagonist sFRP1, which is expressed at the opposite side of Wnt source, accumulates on a novel type of heparan sulfate (HS), N-acetyl-rich HS, which is highly presented in the outer of cardiogenic mesoderm, achieving local inhibition of Wnt signaling by restricting sFRP1 spreading. These two intricate regulatory systems restrict Wnt signaling and ensure reproducible patterning of a thin pericardium tissue.

## Introduction

Morphogens are secreted molecules that pattern embryonic tissues with a concentration gradient that peaks at a localized source and decreases further away. These molecules are important not only for the embryo but also in the adult. However, how the robustness of morphogen distribution is reliably maintained to ensure a reproducible patterning is still debated (Shilo and Barkai, 2017).

The distribution of morphogens is influenced by extracellular molecules. Among them, the receptor is key since it not only transmits the signal into the cell, but can also trap and locally enrich a morphogen on receptor-expressing cells and thereby restrict morphogen distribution on neighboring cells. In addition, many types of receptors mediate internalization of morphogens into cells and subsequent degradation; and/or regulate the morphogen distribution range by ligand-dependent receptor expression. In simulation-based studies, the latter is considered to impart robustness against fluctuations in ligand expression (Eldar et al., 2003). Additionally, the extracellular matrix component heparan sulfate, which binds to morphogens, has an essential role in shaping a morphogen gradient (Yan and Lin, 2009). Secreted antagonists do not only inhibit binding of a ligand to its receptor but also expand the distribution and signaling range of the ligand (Mii and Taira, 2009).

One such morphogen, Wnt, has emerged as a key regulator of vertebrate heart development (Ruiz-Villalba et al., 2016). The so-called canonical Wnt/β-catenin signaling pathway functions to restrict differentiation of myocardium tissue (prospective heart muscle) and promote differentiation of alternative heart tissues, the pericardium (prospective lining of the pericardial cavity). It is still generally unclear which particular *wnt* genes encode relevant Wnt signals to regulate heart development, with possible differences between different vertebrate classes likely, as well as redundancy in some species (Mazzotta et al., 2016; Ruiz-Villalba et al., 2016). It is therefore generally difficult to study the regulation of Wnt signal in vertebrate heart development. However, in *Xenopus* heart development, it has been established that Wnt6 is secreted from the ectoderm-derived epidermis to pattern the adjacent cardiogenic mesoderm in a concentration-dependent manner (Lavery et al., 2008a, 2008b) so that a relatively thin pericardium differentiates close to the epidermis and a broad myocardium further inside the embryo at a distance from the source of Wnt6 ligand (Fig. 1A). However, it is still unclear how Wnt6 protein distribution and the signaling range is regulated to ensure reproducible positioning of pericardium and myocardium in the cardiogenic mesoderm.

**Fig. 1.**
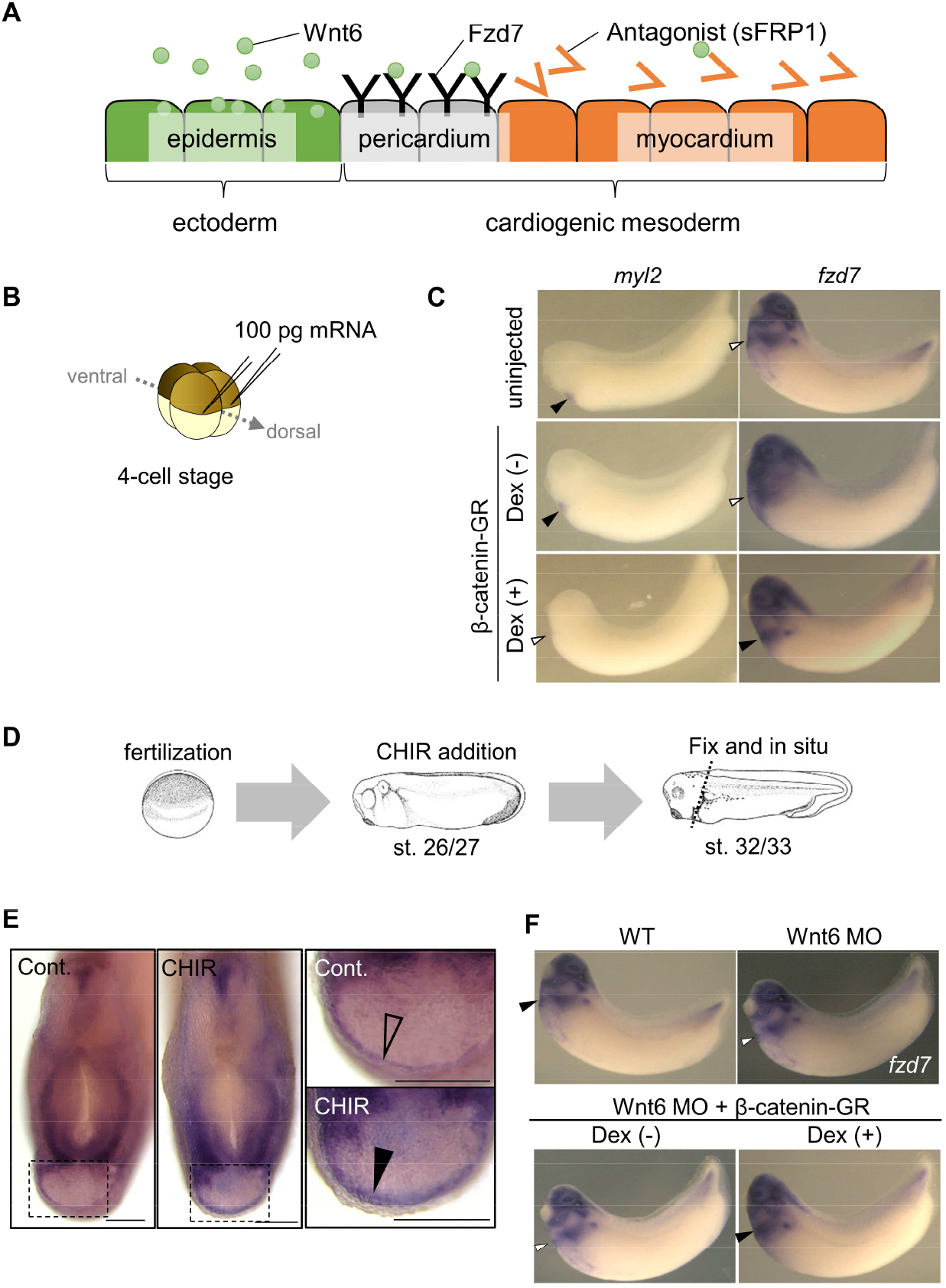
Wnt6/β-catenin signaling induces *fzd7* expression during *Xenopus* heart development. (A) Schematic figure of distributions of Wnt6, Fzd7 and sFRP1 in heart development. Wnt6 is secreted from the epidermis (at the outside of the embryo). The antagonist, sFRP1, is secreted from the prospective myocardium region (at the inside of the embryo). The expression of the Wnt receptor, Fzd7 becomes localized to the pericardium region. The pericardial cavity will subsequently form between the peri- and the myocardium. (B-C) β-catenin activation increased *fzd7* expression. mRNA encoding an inducible β-catenin protein (β-catenin fused with the hormone-binding domain of the human glucocorticoid receptor (GR)) was injected into two dorsal blastomeres at 4-cell stage and the protein was activated at the tailbud stage (st.22-23) with dexamethasone (Dex) (B). β-catenin activation resulted in a decreased in *myl2* expression (n = 17/18) and an increase in *fzd7* expression (n = 18/19) around the heart region (arrowhead, Dex (+)), compared with the control (Dex (-); n = 22/23, n = 18/20, respectively) (C). (D-E) Embryos were treated with a Wnt agonist, CHIR99021 (5 μM; control DMSO) from st. 26/27, which is just before the onset of Wnt6 expression, to st. 32/33 (D). *fzd7* expression area was broader with CHIR treatment (arrowhead; n = 30/36), but not with DMSO control (open arrowhead; n = 1/23). (F) Wnt6 MO was injected into the animal pole at one-cell stage alone or together with mRNA of inducible β-catenin-GR (as indicated in Supplemental Fig. 1E). β-catenin-GR was activated at the tailbud stage with Dex (where indicated by Dex (+)). Note that *fzd7* expression decreased by *wnt6* knockdown (n = 17/18 (Wnt6 MO), n = 17/17 (Wnt6 MO + β-catenin-GR (Dex (-))) and was rescued by experimental β-catenin activation (n = 18/20). See Suppl. Fig. 1F-G for concomitant reduction of myocardial marker expression (*myl2*). Scale bar = 200 μm.

In the early embryo, the range of Wnt8 signal is precisely regulated by two novel types of heparan sulfates (HS), N-sulfo-rich HS and N-acetyl-rich HS, and secreted Wnt binding proteins including Frzb (also known as sFRP3) (Mii et al., 2017; Mii and Taira, 2009). Wnt8 distribution range can be shortened by its binding to the receptor and to N-sulfo-rich HS. Frzb/sFRP3, originally described as a secreted Wnt antagonist, and its binding partner N-acetyl-rich HS prevents Wnt8 from binding to its receptor and N-sulfo-rich HS, making Wnt8 distribution range longer. In heart tissue, sFRP1, which can function as a secreted antagonist of Wnt6, is expressed in the prospective myocardium region (Gibb et al., 2013; Xu et al., 1998). We wondered whether similar mechanisms as in the early embryo may operate to regulate the distribution of Wnt6 in the cardiogenic mesoderm.

The Wnt receptor Frizzled7 (Fzd7) is expressed in the cardiogenic mesoderm, which is essential for heart development (Abu-Elmagd et al., 2017; Wheeler and Hoppler, 1999). The expression of *fzd7* is known to be increased by Wnt signaling in *Xenopus* neuroectoderm and human embryonic carcinoma cells (Willert et al., 2002; Young et al., 2014), but there are no such reports in heart development.

In this study, we analyzed the regulatory mechanism involving Wnt signaling in cardiogenic mesoderm differentiation ensuring the reproducible patterning of a thin tissue, pericardium, especially focusing on the receptor expression, the secreted antagonist sFRP1 and HS.

## Results

### *fzd7* expression is regulated by Wnt6 signaling

As the cardiogenic mesoderm becomes patterned into peri- and myocardium, initially broad *fzd7* expression changes into being maintained just in the prospective pericardium (Wheeler and Hoppler, 1999; Supplemental Fig. 1A), while little expression remains in the prospective myocardium. This pattern of expression reminded us of the expression pattern of the pericardium marker *gata5* in early heart development, which is positively regulated by Wnt signaling (Gibb et al., 2013). We therefore wondered whether the expression of *fzd7* was regulated by Wnt signaling in heart development.

To test whether Wnt6 signaling is capable of regulating *fzd7* expression, we injected mRNA encoding an inducible β-catenin (Glucocorticoid Receptor (GR)-fused β-catenin) (Afouda et al., 2008) at two-cell stage and activated it at the tailbud stage (st. 22-23) by dexamethasone (Dex) treatment (Fig. 1B). The expression of *fzd7* at stage 32/33 increased throughout the heart region (Fig. 1C, β-catenin-GR + Dex (+), arrowhead), thereby also indicating expansion of pericardium region. In this condition, the expression of a myocardium marker gene, *myosin light chain 2* (*myl2*) decreased probably due to concomitant reduction of myocardium tissue. This Wnt-dependent *fzd7* expression was confirmed by targeted overexpression of Wnt6 by DNA injection (Supplemental Fig. 1B-D). As further confirmation, we treated embryos at the relevant stage of development with a Wnt signaling agonist, CHIR99021 (Fig. 1D). We found *fzd7* expression expanding throughout what appears to be the cardiogenic mesoderm (Fig. 1E). These results are also consistent with previous reports that overexpression of the Wnt antagonist sFRP1 diminishes the expression of *fzd7* in the prospective pericardium, and *sfrp1* knockdown increases *fzd7* expression (Gibb et al., 2013). In addition, experimental knockdown of endogenous *wnt6* (by injection of antisense morpholino oligonucleotide (MO) for *wnt6* (Lavery et al., 2008a) at one-cell stage) subsequently reduced *fzd7* expression around the heart region; and remarkably, *fzd7* expression was reinstated by β-catenin activation at the tailbud stage (Fig. 1F). Taken together, these results show that *fzd7* is expressed in heart development in a Wnt-dependent manner.

### Fzd7 restricts range of Wnt6 protein distribution

Wnt6 is expressed in the epidermis overlaying the developing embryonic heart organ (Lavery et al., 2008a) and functions to restrict myocardial and promote pericardial tissue differentiation (Gibb et al., 2013; Lavery et al., 2008b), which results in patterning of the cardiogenic mesoderm into prospective myocardial tissue only at a distance from Wnt6 secreting cells and pericardial differentiation nearby (Fig. 1A). To examine whether Fzd7 restricts Wn6 protein distribution, we examined distribution of a Wnt6 protein with an mVenus-tag (mV-Wnt6). The functional activity of N-terminus tagged Wnt6 (mV-Wnt6) was nearly the same as intact Wnt6 (Supplemental Fig. 2). At 4-cell stage, we injected mRNA of mV-Wnt6 and Fzd7 (with mRFP as a tracer) into two different blastomeres (Fig. 2A). mV-Wnt6 protein was highly accumulated on Fzd7-expressing cells (Fig. 2B, C). These results indicate that Fzd7 accumulates Wnt6 on the cell membrane.

**Fig. 2.**
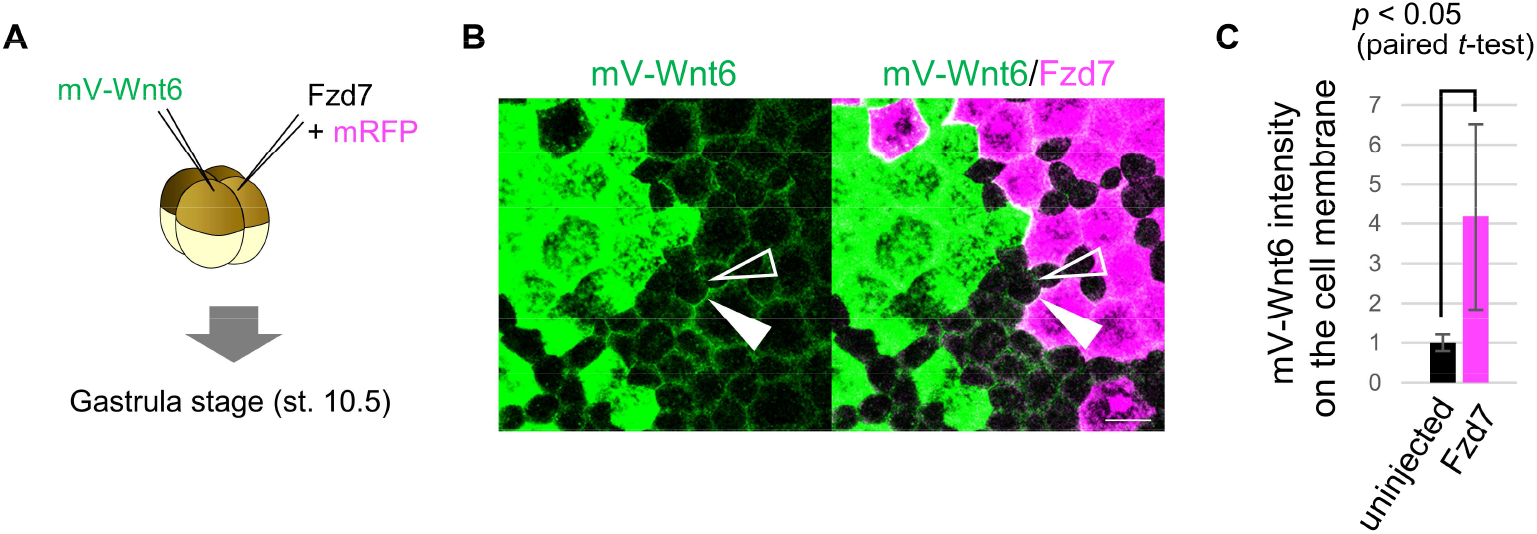
Wnt6 distribution is restricted by Fzd7. (A) Schematic view of the experimental process. 500 pg of mV-Wnt6 mRNA and Fzd7 mRNA (with mRFP) were injected into different blastomere at 4-cell stage, and the specimens were fixed at st.10.5 (gastrula stage). (B) mV-Wnt6 accumulates on Fzd7-expressing cells (confocal image). mV-Wnt6 (green) was accumulated on Fzd7-overexpressed cells (magenta; arrowhead), not on intact cells (not Fzd7-overexpressed cells, open-arrowhead). Scale bar, 30 μm. (C) Signal intensity of mV-Wnt6 with/without Fzd7 expression (n = 4, 6, respectively). Bars represent SD.

### Exploring functional significance of feedback regulation

To explore the potential biological significance of the Wnt-dependent expression of *fzd7* in heart development, we examine it with mathematical modeling.

Model simulations indicate that Wnt-dependent expression of *fzd7* functions to restrict and enhance Wnt-signal activity around the source of Wnt6 (Fig. 3A). Considering a threshold shown in the graph (Fig. 3A, merge), the feedback regulation is suggested to be enough for activation of Wnt signaling in a well-defined narrow band. However, we wondered if such a narrow patterning can be achieved without this feedback regulation but instead hypothetically with an initially high expression of the receptor in the prospective pericardium region because receptors can restrict the distribution of ligands. To examine it, we assumed various initial amounts of the receptor. As a result, the high expression in the initial condition makes a steep gradient (Supplemental Fig. 3; for instance, feedback: 0, initial amount: 8). Thus, such narrow-band Wnt signaling activation can be theoretically achieved with either condition.

**Fig. 3.**
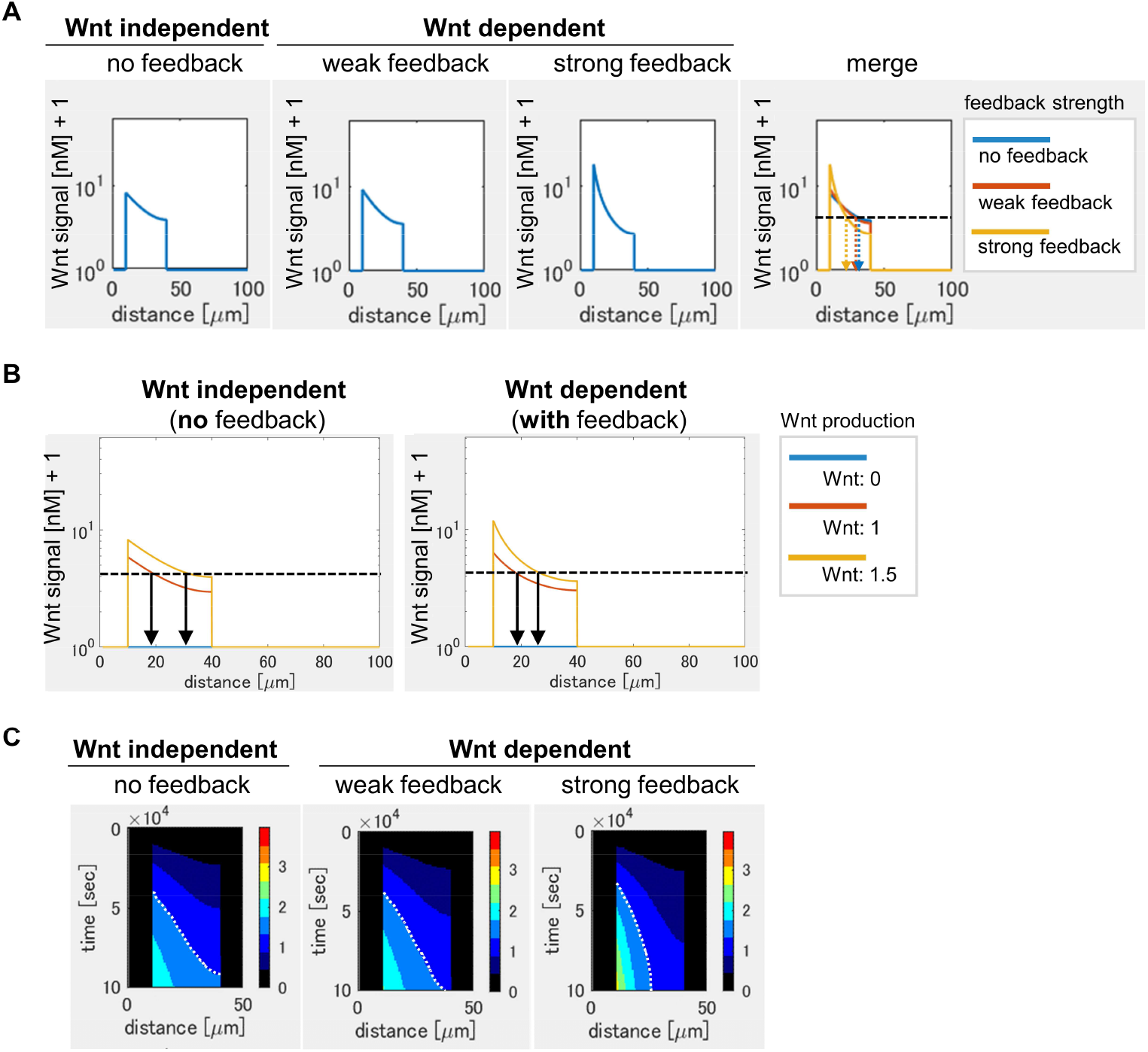
Feedback regulation of Fzd7 expression restricts Wnt signaling range in a mathematical model. (A) Wnt-dependent expression of Fzd7 increases and restricts Wnt signaling, making a steep gradient. An example of the threshold was shown by a dashed line (merge). The arrows showed the edge of the activated region in each condition. The “strong feedback” means 5-fold strength of the feedback compared with the “weak feedback”. These figures correspond to Supplemental Fig. 3 (initial amount: 2). (B) The feedback regulation of *fzd7* expression makes Wnt gradient robust to Wnt production fluctuation. One example of a threshold was shown in a dotted line. Arrows indicated the edge of Wnt-signal activated region in the threshold. Relative rates of Wnt production were shown in the right. The figures of “Wnt independent” and “Wnt dependent” correspond to Supplemental Fig. 4 (feedback: 0, initial amount: 2; feedback: 5, initial amount 1, respectively). (C) Heatmap image of Wnt activity. The time change was shown as kymographs, in which time proceeded from the upper to the bottom. Dotted line in the graph shows an example of temporal changes in a Wnt-activated region. The line was a nearly vertical line in the strong feedback condition, indicating that the system with a strong feedback can quickly reach a steady state. These figures correspond to Supplemental Fig. 5C (the initial amount of the receptor: 2).

Developmental patterning in different contexts is generally challenged by fluctuation of ligand production. To further examine the importance of the feedback, we focused on fluctuation of ligand production. It has been suggested that ligand-dependent expression of the receptor endows the system to become robust to ligand production fluctuations (Eldar et al., 2003). They simulated the signal activation only in a steady state. However, in most in vivo situations, it is not always possible to assume a steady state. Actually, it has been noted that extracellular factors, such as HS proteoglycan (HSPG), delay the time to reach a steady state. So, we challenged our model with 50% fluctuation in the ligand production in a simulation not limited to a steady state (Fig. 3B, Supplemental Fig. 4A). We found that the changes in boundary position (arrows in Fig. 3B) in response to the Wnt production change seem to be smaller with the feedback than that without feedback. To further confirm it, we calculated the differences of boundary position between the two Wnt production rates (Supplemental Fig. 4B) and the differences were found to be small in all thresholds with the feedback and/or an excess amount of initial receptor. These mathematical simulations suggest that the system with the feedback regulation is robust to the fluctuation, not only at a steady state, but also in a transient state. In addition, an excess amount of initial receptor also makes the gradient robust to the fluctuation.

In the above simulation results, the signal levels were shown at a certain time point (∼ 1 day after the onset of the simulation), which corresponds to the time for heart development in *Xenopus*. Developmental systems are, however, also often challenged by variation in speed. For a more detailed examination, we next visualized the time change of the activation level (and the amounts of free receptor and free ligands) by a heat map (Fig. 3C, Supplemental Fig. 5A-C). The dotted line aligned between two colors was almost vertical with the feedback, but not with no feedback (Fig. 3C), meaning that the system with the feedback quickly reaches a steady state in Wnt signaling. These mathematical simulations indicate that feedback is robust not only to changes in Wnt secretion level but also to changes in a developmental time window.

We next examined how Wnt-dependent expression of *fzd7* might be required for Wnt signaling activation only in a narrow region in vivo. To examine this, we synthesized a plasmid construct that expresses Fzd7 lacking the intracellular domain required for Wnt-signaling that was fused with a transmembrane sequence by modification of the TOPFLASH plasmid (M&M, Supplemental Fig. 6A). We expected that this construct would cell-autonomously inhibit Wnt signaling/Fzd7 expression in a Wnt-dependent manner. We confirmed that the injection of this construct increases Wnt ligand on the cell membrane but inhibits Wnt signaling when Wnt6 was overexpressed at gastrula stage (Supplemental Fig. 6B, C). When the plasmid was injected into the prospective heart region (Supplemental Fig. 6D), this injection cell-autonomously decreased *fzd7* expression as expected in the injected cells also during the heart developing stage (st. 32) but, interestingly, it also expanded *fzd7*-expression into the prospective myocardium region (Supplemental Fig. 6E). Thus, it suggests that Wnt-dependent feedback expression of Fzd7 is required to maintain Wnt-signal activation in a thin region.

### sFRP1 expands range of Wnt6 distribution

This Wnt6 and Fzd7 feedback regulation operates in a wider context of known regulation by sFRP1 and likely regulation by heparan sulfate. sFRP1 is a secreted antagonist of Wnt ligand, and is essential for normal myocardium differentiation (Gibb et al., 2013). We therefore initially expected that sFRP1 inhibits Wnt6 signaling. However, in early *Xenopus* development, a related protein, Frzb (aka Sfrp3), was found to expands the range of Wnt8 distribution (Mii and Taira, 2009). So, we examined whether sFRP1 functions to expand Wnt6 protein distribution. To analyze each distribution of Wnt6 and sFRP1, we used mVenus-tagged proteins and revealed that Wnt6 seems to have a narrow distribution range staying relatively close to Wnt6-secreting cells whereas the distribution range of sFRP1 seems to be broad (Supplemental Fig. 7A).

Then, we next analyzed whether sFRP1 can expand the distribution range of Wnt6. The distribution of Wnt6 was expanded by sFRP1 (Supplemental Fig. 7A). Although in this experiment, Wnt6 and sFRP1 were co-injected into the same blastomere, the sources of these molecules are different in vivo. So, we next injected them into different blastomeres. Consistent with the above experiment, it was confirmed that the distribution range of Wnt6 was wider when sFRP1 was injected (Supplemental Fig. 7B).

These results indicate that the antagonist sFRP1 can expand the Wnt6 ligand distribution, maybe preventing its binding to the Fzd7 receptor, similar to the case of Wnt8-Frzb (Mii et al., 2017).

### Involvement of heparan sulfate in the heart development

In general, morphogen distribution is considered to be regulated by its binding to heparan sulfate (Yan and Lin, 2009). Two types of HSPG modification, N-acetyl and N-sulfo, are found to be involved in Wnt8-mediated signaling (Mii et al., 2017). To examine the involvement of two types of heparan sulfate in the Wnt6-mediated signaling, we utilized experimental expression of Ndst1 enzyme, which converts N-acetyl HS to N-sulfo HS. Although Wnt6 distribution was not substantially changed by Ndst1 expression (Fig. 4A), sFRP1 distribution was changed, and could not be detected on the Ndst1-expressing cells, suggesting that sFRP1 preferentially localizes to N-acetyl HS (Fig. 4A). Consistently, sFRP1 was preferentially localized to *ndst1*-knockdowned cells (Supplemental Fig. 7C). To further confirm it, we visualized N-acetyl-rich HS and N-sulfo-rich HS by immunohistochemistry (IHC). sFRP1 was found to be highly colocalized with N-acetyl-rich HS (Fig. 4B), not with N-sulfo-rich HS (Supplemental Fig. 7D). In the heart, N-acetyl-rich HS was clearly present in the outer of cardiogenic mesoderm, while N-sulfo-rich HS was not clearly detected in the heart region (Fig. 4C). Consistent with this, endogenous *ndst1* expression was not clearly observed in the heart region (data not shown).

**Fig. 4.**
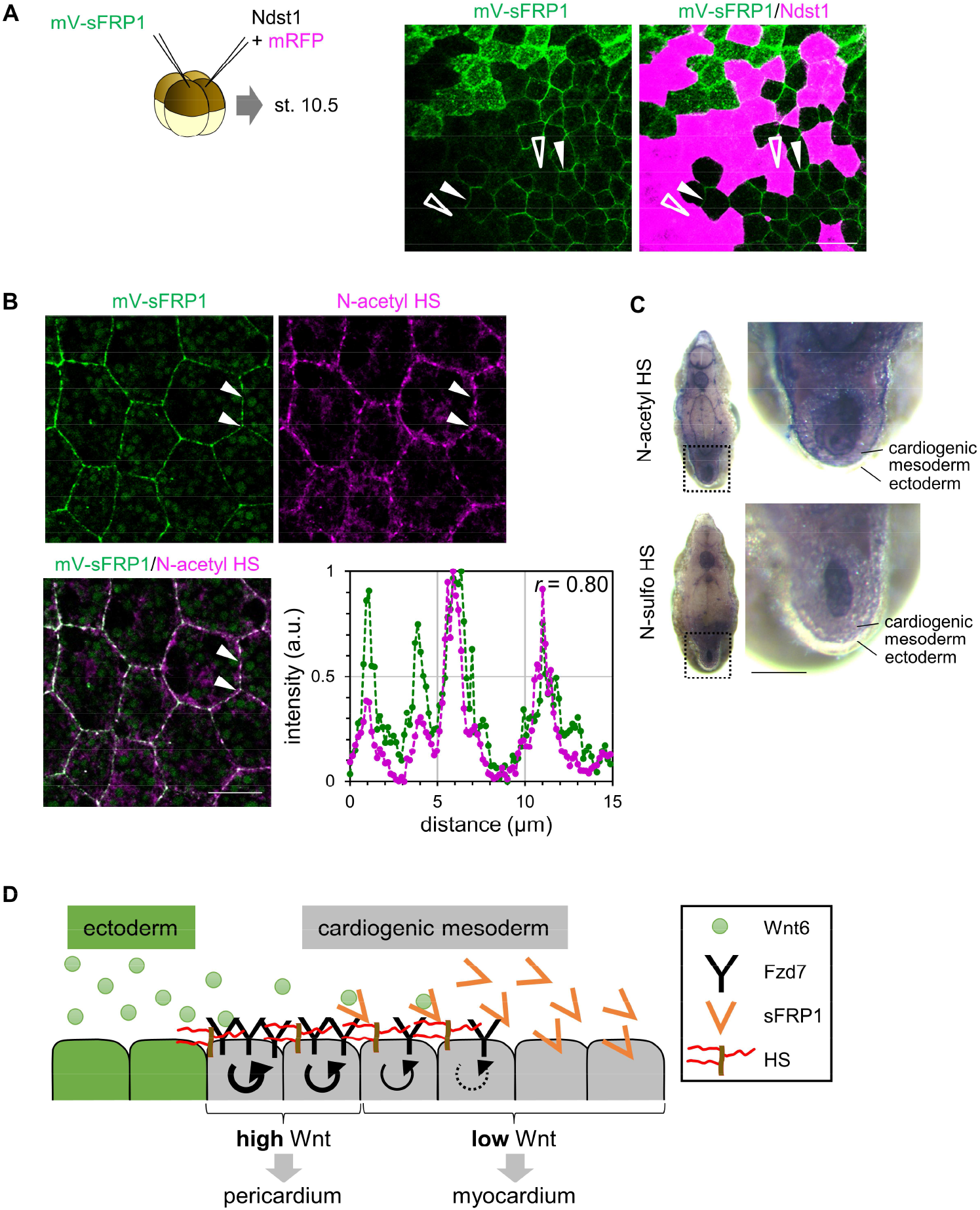
Heparan sulfate in *Xenopus* heart development. (A) sFRP1 did not accumulated on *ndst1*-expressing cells (open arrowhead), compared to the intact cells (arrowhead). Indicated mRNAs were injected into different blastomeres at 4-cell stage. *ndst1* was expressed in the magenta-colored cells. Scale bar, 30 μm. (B) sFRP1 was highly localized on N-acetyl-rich HS. mV-sFRP1 mRNA was injected into one blastomere at 4-cell stage. N-acetyl-rich HS was visualized by IHC. Signal intensity was measured between two arrowheads (green, mV-sFRP1; magenta, N-acetyl-rich HS). Correlation coefficient (*r*) was shown in the graph. Scale bar, 20 μm. (C) The distribution of N-acetyl-rich and N-sulfo-rich HS in the heart tissue. IHC for N-acetyl-rich and N-sulfo-rich HS. The dotted line in the left figure shows the position of magnified image on the right. N-acetyl-rich HS was highly localized in the outer of cardiogenic mesoderm. Scale bar, 0.1 mm. (D) Schematic view of Wnt signaling in the heart tissue. Wnt ligand is secreted from the epidermis (left cells). Before the onset of Wnt6 expression, the receptor (Fzd7) is broadly expressed around the prospective pericardium region. Wnt signal is activated in the concentration dependent manner (circular arrows), which induces Fzd7 expression and restricts Wnt ligand spreading. sFRP1 is secreted from the prospective myocardium region. This distribution is potentially longer than that of Wnt6, but is restricted by N-acetyl HS, which is highly expressed at the outer of cardiogenic mesoderm.

As shown above, sFRP1 potentially expands Wnt6 distribution and shows longer distribution, compared with Wnt6. However, extracellular sFRP1, as it is secreted from the prospective myocardial region, may be trapped by N-acetyl-rich HS, and thereby inhibits Wnt signaling just around the myocardium region. This will contribute to making the Wnt-activated region narrow (Fig. 4D).

Consistently, theoretical analysis showed that the Wnt-activated region will as a consequence become narrow with sFRP1- and HS-mediated mechanisms (Supplemental Fig. 5D), compared with no sFRP1/HS condition (Supplemental Fig. 5C). In addition, lines between two colors seem to become vertical faster with sFRP1(Supplemental Fig. 5C, arrows), suggesting that the system reaches a steady state in Wnt signaling faster with sFRP1/HS.

## Discussion

In this study, through analyzing patterning of the cardiogenic mesoderm to specify a normally thin pericardium, where endogenous Wnt signaling is highly activated, we revealed that the Wnt receptor Fzd7 is expressed in a Wnt-dependent manner, and that this feedback facilitates robust formation of a steep gradient of Wnt ligand distribution and thus Wnt pathway activation. In addition, sFRP1 preferentially localizes on N-acetyl-rich HS, which is highly presented in the outer of cardiogenic mesoderm, implying that N-acetyl HS traps sFRP1, thereby not allowing sFRP1 to expand much further into the prospective pericardial region, contributing to shaping a steep gradient of Wnt activation and keeping the future pericardium thin (Fig. 4D).

To explore the functional significance of feedback regulation we used mathematical modeling. To restrict a highly activated region in morphogen-mediated signaling, it was theoretically feasible to form such a narrow activation when an excess number of receptors are initially present, even without this feedback regulation (Supplemental Fig. 3). However, in terms of energy used to produce high levels of Fzd proteins throughout the cardiogenic mesoderm to ensure patterning of a thin pericardium, this theoretical strategy may not be compatible with Darwinian evolution: maybe starting with a low expression everywhere and then specifically maintaining higher levels where necessary, through the discovered natural feedback mechanisms, may be more compatible with achieving higher fitness in the ‘wild’. Since ∼83% of genes are differentially expressed among individuals in humans (Storey et al., 2007), the production of Wnt ligand is also expected to fluctuate. In this case (with Wnt fluctuation), the system with the feedback was more robust to achieve reproducible patterning. Therefore, the feedback that we discover is key to ensuring efficient and reliable patterning. There are many things to perturb the reproducibility of morphogen-mediated signaling gradients in vivo. For instance, the speed of morphogenesis can be variable, e.g. by temperature (i.e., in fishes and frogs). Therefore, the amount of Wnt ligand, and the length of time for the heart differentiation can be expected to fluctuate among individuals in vivo. At least in these two situations, the discovered feedback regulation is found to ensure the reproducibility as shown above.

The knockdown of sFRP1 expands the pericardium region (Gibb et al., 2013), suggesting that the feedback on *fzd7* expression is not sufficient to ensure “thin” patterning. However, when we experimentally tried to interfere with this feedback, we find this expands the pericardium region in vivo (Supplemental Fig. 6D), suggesting that the *fzd7* feedback contributes towards, and is necessary for, normal patterning. In addition, the time to reach a steady state in Wnt signaling was faster with sFRP1 and heparan sulfate (Supplemental Fig. 5D). These results suggest that together these two regulatory mechanisms, the *fzd7* feedback and sFRP1 function with N-acetyl HS, ensure reproducible thin patterning of the pericardium.

We revealed that a modification of heparan sulfate, N-acetyl HS, is present in the outer of cardiogenic mesoderm. In addition, N-acetyl modification of HS had previously been considered to be just a precursor of other modifications, but sFRP1 was found to be one of the specific binding partner of N-acetyl HS in addition to Frzb (Mii et al., 2017) and other proteins related to morphogens (Yamamoto, in preparation). These results imply that the difference in the preference of N-acetyl binding possibly determines the distribution/signaling ranges of ligands in general. Therefore, understanding of each function of HS modifications including the “precursor” is important.

Wnt6 and sFRP1 molecules not only regulate normal embryonic heart development, but also regulate repair and regeneration after heart muscle injury in animal models of heart attack (myocardial infarction). Our results will provide important knowledge that is likely going to be relevant for medical applications, for instance for drug design since cell-surface molecules that we study, such as the Frizzled, or a specific modification of heparan sulfate, or even the secreted molecule sFRP1, generally provide better drug targets than molecules inside cells.

It should be noted that we still do not know how much cell division and cell movement there are in the cardiogenic mesoderm while Wnt-mediated patterning is going on, but they should be analyzed to understand the mechanism of the reproducibility in the heart development. Possibly, Fzd7 feedback and sFRP1 function could also be important for providing robustness for that.

There are many extracellular binding partners of morphogens, including the receptors, the antagonists and heparan sulfates, which can change morphogen distribution. To reveal the precise regulation of morphogens and consider the medical applications, the regulatory mechanisms of these components must be investigated further.

## Material and Methods

### *Xenopus* embryo manipulation and microinjection

All animal experiments were approved by The Office for Life Science Research Ethics and Safety, the University of Tokyo. For experiments conducted at the University of Aberdeen, all animal experiments were carried out under license from the United Kingdom Home Office (PPL PA66BEC8D). Manipulation of *Xenopus* embryos and microinjection experiments were carried out according to standard methods as previously described (Sive et al., 2000). Briefly, unfertilized eggs were obtained from female frogs injected with gonadotropin, and artificially fertilized with testis homogenate. Fertilized eggs were dejellied with 2% L-cysteine-HCl solution (pH7.8), and incubated in 1/10x Steinberg’s solution at 14-20ºC. Embryos were staged according to Nieuwkoop and Faber (Nieuwkoop and Faber, 1967). The amounts of injected mRNAs are described in the figure legends. For the experiments with the Wnt/β-catenin signaling agonist CHIR-99021, embryos were left to develop in 0.1×Steinberg’s solution to embryonic stage 26, and incubated in 5 μM solution (or DMSO as a control).

### Plasmid and RNA construction

The insertion sequence for the plasmids of monomeric Venus (mVenus/mV) fused with Wnt6 or sFRP1 were subcloned from pCS2+xWnt6 (Lavery et al., 2008b) or pXT7-xFrzA (a gift from Dr. Sergei Sokol) (Xu et al., 1998), respectively. The PCR products for N-terminus tagged construct were inserted into pCSf107SP-mV-mT, and those for C-terminus tag were into pCSf107-mcs-mV-mT. An inducible β-catenin (Glucocorticoid Receptor (GR)-fused β-catenin) was used as previously described (Afouda et al., 2008). *Fzd7* fragment that was cut out from pCS2+xFz7-ASN (a gift from Dr. Masanori Taira) was inserted into pCS2+mcs-6MT-T vector using *Eco*RI/*Age*I sites.

TCF-dnFrizzled7-transmembrane plasmid (TCF-dnFzd7-TM) was a modified plasmid from the TOPFLASH system, which is a well-known Wnt reporter construct, expresses luciferase in a Wnt-dependent manner. We replaced the luciferase sequence by the extracellular domain of *fzd7* (dominant negative Fzd7, dnFzd7) with mouse IgG transmembrane sequence (TM) (Supplemental Fig. 6A). This construct expresses the extracellular domain of Fzd7 on the cell surface dependent on the Wnt signaling, which cell-autonomously inhibits Wnt-dependent *fzd7* expression.

mRNAs were transcribed in vitro using mMessage mMachine SP6 kit (Ambion). All the primers for clonings were listed in Supplemental material Table S2.

### qRT-PCR

Total RNA was isolated from whole embryos using the RNeasy Mini Kit, according to manufacturer’s instructions (QIAGEN) for processing of animal tissues (see also Lee-Liu et al., 2012; Nakamura et al., 2016). The abundance of RNAs was determined using a LightCycler 480 and SYBR Green I Master Reagents (Roche). Relative expression levels of genes were determined using ΔΔCt.

### Whole mount in situ hybridization (WISH)

WISH was performed based on *Xenopus* standard methods (Harland, 1991) with slight modifications in the duration of washes with a hybridization-temperature of 65°C. The plasmid for the probes were linearized and transcribed in vitro using DIG RNA labeling mix (Roche; supplementary material Table S1 for plasmid templates for RNA synthesis).

### Immunohistochemistry

*Xenopus* gastrula embryos were fixed with MEMFA (0.1 M MOPS, pH 7.4, 2 mM EGTA, 1 mM MgSO4, 3.7% formaldehyde) and immunostained by standard protocols with Tris-buffered saline (Sive et al., 2000). The specimens were incubated with the following primary antibodies overnight at 4ºC: anti-N-acetyl HS (NAH46 (Suzuki et al., 2008), in-house preparation, 1:50), anti-N-sulfo HS (HepSS-1 (Kure and Yoshie, 1986), in-house preparation, 1:1000), diluted with 2%BSA in TBT (0.01% TritonX-100 in TBS). Following this, the samples were incubated with the secondary antibodies overnight at 4ºC: anti-rabbit or mouse Alexa 488 or 555 antibody (Invitrogen).

### Theoretical Model

We used one-dimensional reaction-diffusion equations to simulate the Wnt-signal gradient formation. We assumed that two molecules of Fzd7 bind one molecule of Wnt based on recent X-ray structural analysis (Hirai et al., 2019).

## Abbreviations

W: Wnt6
S: sFRP1
WS: Wnt6-sFRP1
H: Heparan sulfate
SH: sFRP1-Heparan sulfate
WSH: Wnt6-sFRP1-Heparan sulfate
R: Fzd7
WR_2_: Wnt6-Fzd7

Assumptions:

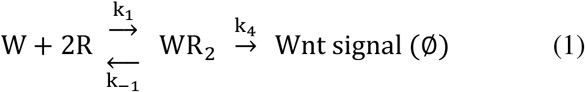

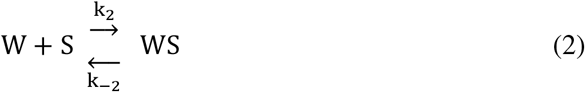

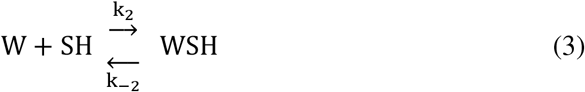

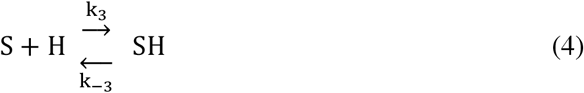

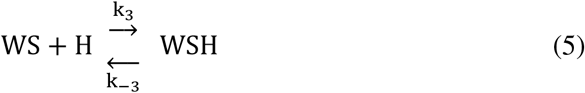

The concentration of molecules follows reaction-diffusion equations below.

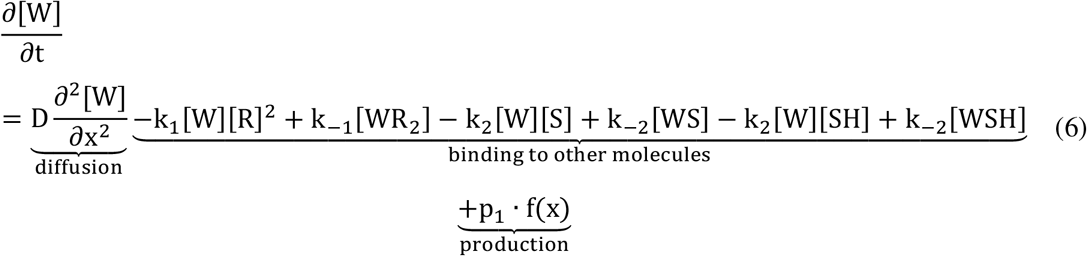

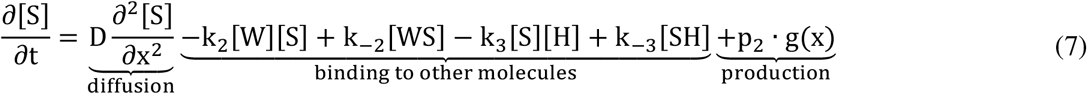

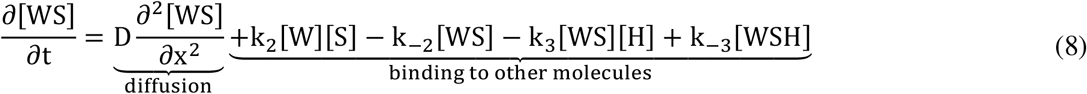

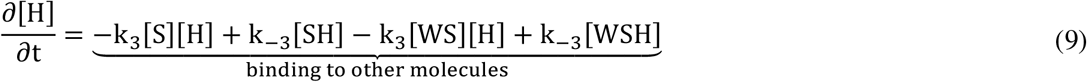

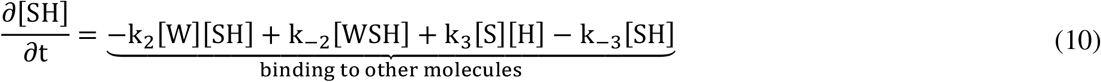

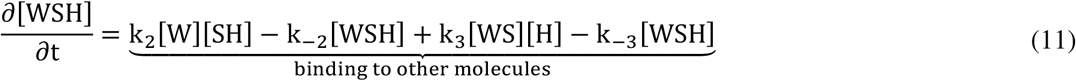

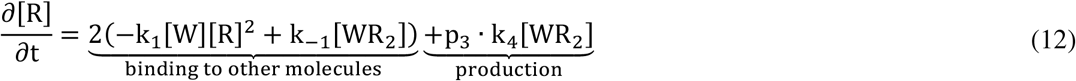

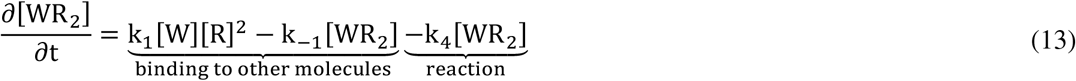

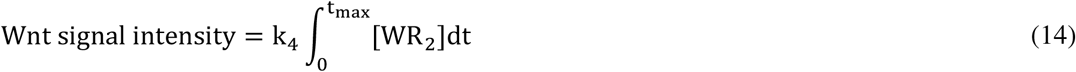

Wnt signal intensity is defined as the integration of the endocytosis rate of Wnt6-receptor complex (equation (14), t_max_ = 10^5^ seconds ≒ 1 day).

Production of Wnt6 or sFRP1 were limited to left or right regions, respectively.

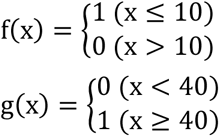

The initial concentrations of the molecules are set to be zero, except for that of the receptor (R_0_) and Heparan sulfate (H_0_). Initially, the receptor is assumed to be expressed in 10 < x < 40 at low level.

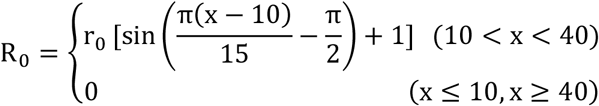

Since Wnt6, sFRP1 and Wnt6-sFRP1 complex is impermeable at the distal end of the sources, the Neumann boundary condition is assumed (15).

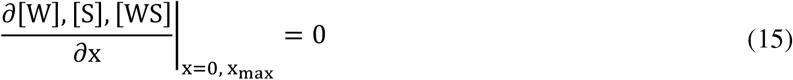

Numerical simulation was performed using the partial differential equation solver (pdepe) in MATLAB (MathWorks, version: R2020a).

## Parameter Values

Parameter values that were needed in our simulation were not reported enough in Wnt signaling. In general, these values do not vary at the order level between morphogens, so we used mainly BMP values as follows.

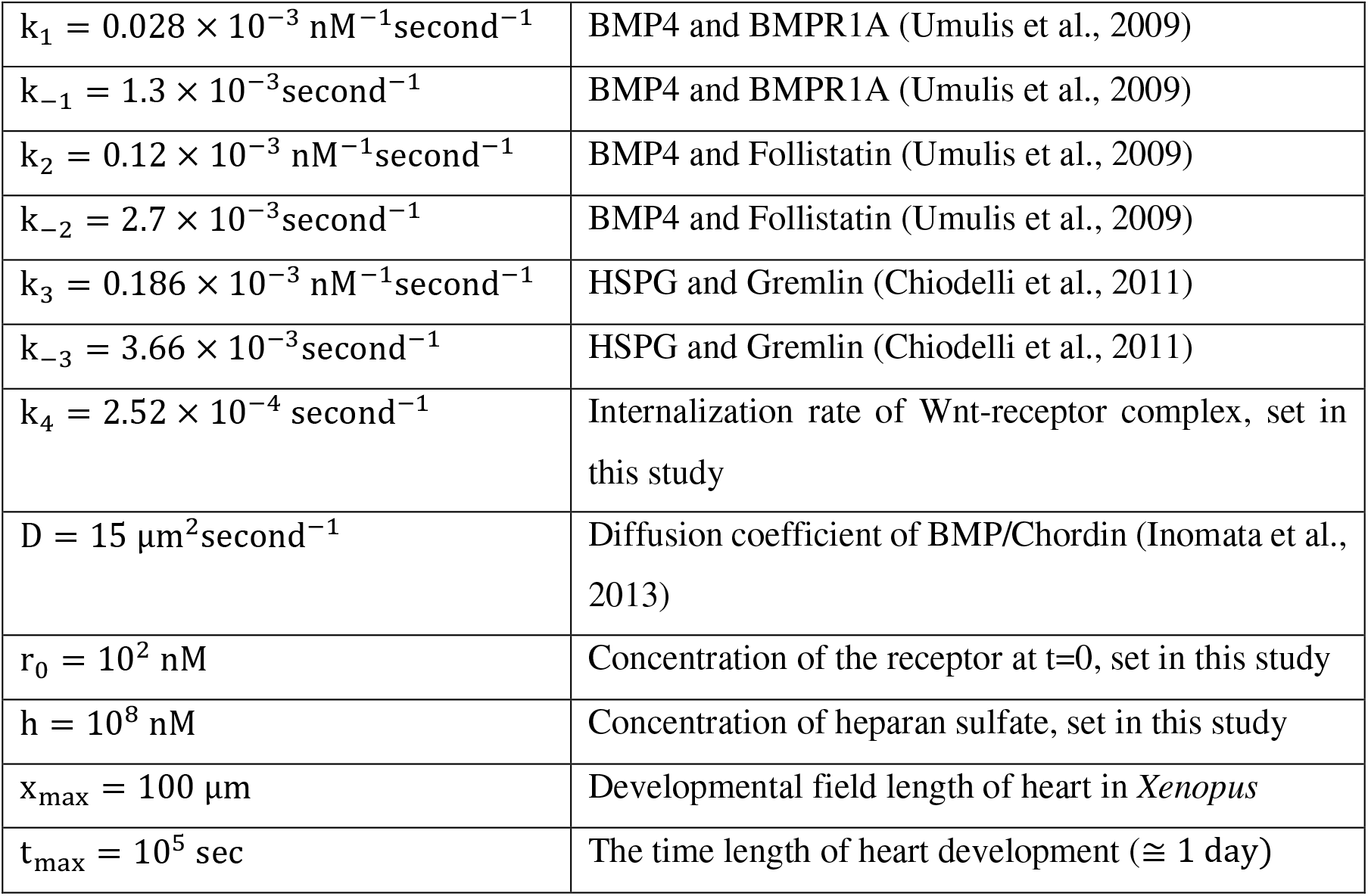

|  |  |
| --- | --- |
| $k_1 = 0.028 \times 10^{-3} \text{ nM}^{-1}\text{second}^{-1}$ | BMP4 and BMPR1A (Umulis et al., 2009) |
| $k_{-1} = 1.3 \times 10^{-3}\text{second}^{-1}$ | BMP4 and BMPR1A (Umulis et al., 2009) |
| $k_2 = 0.12 \times 10^{-3} \text{ nM}^{-1}\text{second}^{-1}$ | BMP4 and Follistatin (Umulis et al., 2009) |
| $k_{-2} = 2.7 \times 10^{-3}\text{second}^{-1}$ | BMP4 and Follistatin (Umulis et al., 2009) |
| $k_3 = 0.186 \times 10^{-3} \text{ nM}^{-1}\text{second}^{-1}$ | HSPG and Gremlin (Chiodelli et al., 2011) |
| $k_{-3} = 3.66 \times 10^{-3}\text{second}^{-1}$ | HSPG and Gremlin (Chiodelli et al., 2011) |
| $k_4 = 2.52 \times 10^{-4} \text{ second}^{-1}$ | Internalization rate of Wnt-receptor complex, set in this study |
| $D = 15 \mu\text{m}^2\text{second}^{-1}$ | Diffusion coefficient of BMP/Chordin (Inomata et al., 2013) |
| $r_0 = 10^2 \text{ nM}$ | Concentration of the receptor at $t=0$, set in this study |
| $h = 10^8 \text{ nM}$ | Concentration of heparan sulfate, set in this study |
| $x_{\text{max}} = 100 \mu\text{m}$ | Developmental field length of heart in *Xenopus* |
| $t_{\text{max}} = 10^5 \text{ sec}$ | The time length of heart development ($\cong 1 \text{ day}$) |

We non-dimensionalized the reaction diffusion equation with the following scaling factors:

4 We non-dimensionalized the reaction diffusion equation with the following scaling factors:
|  |  |
| --- | --- |
| $t^* = t_{\text{max}} - t_{\text{min}}$ | scaling factor for non-dimentionalization of time |
| $x^* = \sqrt{4D(t_{\text{max}} - t_{\text{min}})}$ | scaling factor for non-dimentionalization of distance |
| $c^* = p_1 \cdot (t_{\text{max}} - t_{\text{min}})$ | scaling factor for non-dimentionalization of concentration |

## Nondimensional Equations

Using these parameters, we non-dimensionalized parameters and variables in the reaction diffusion equations as follows:

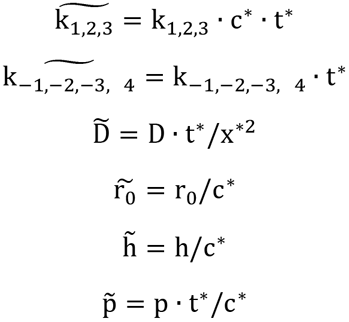

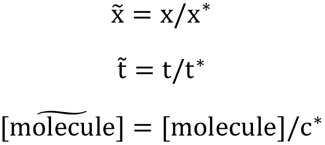

**Supplemental material Table S1.** Summary of antisense RNA constructs for probe synthesis.

| Gene | Restriction enzyme | Transcription Enzyme | References |
| --- | --- | --- | --- |
| *fzd7* (Full-length) | *HindIII* | T3 | This paper |
| *fzd7* (intracellular domain) | *AvaI* | SP6 | This paper |
| *myl2* | *BamHI* | T7 | (Evans et al., 1995) |

**Supplemental material Table S2.**
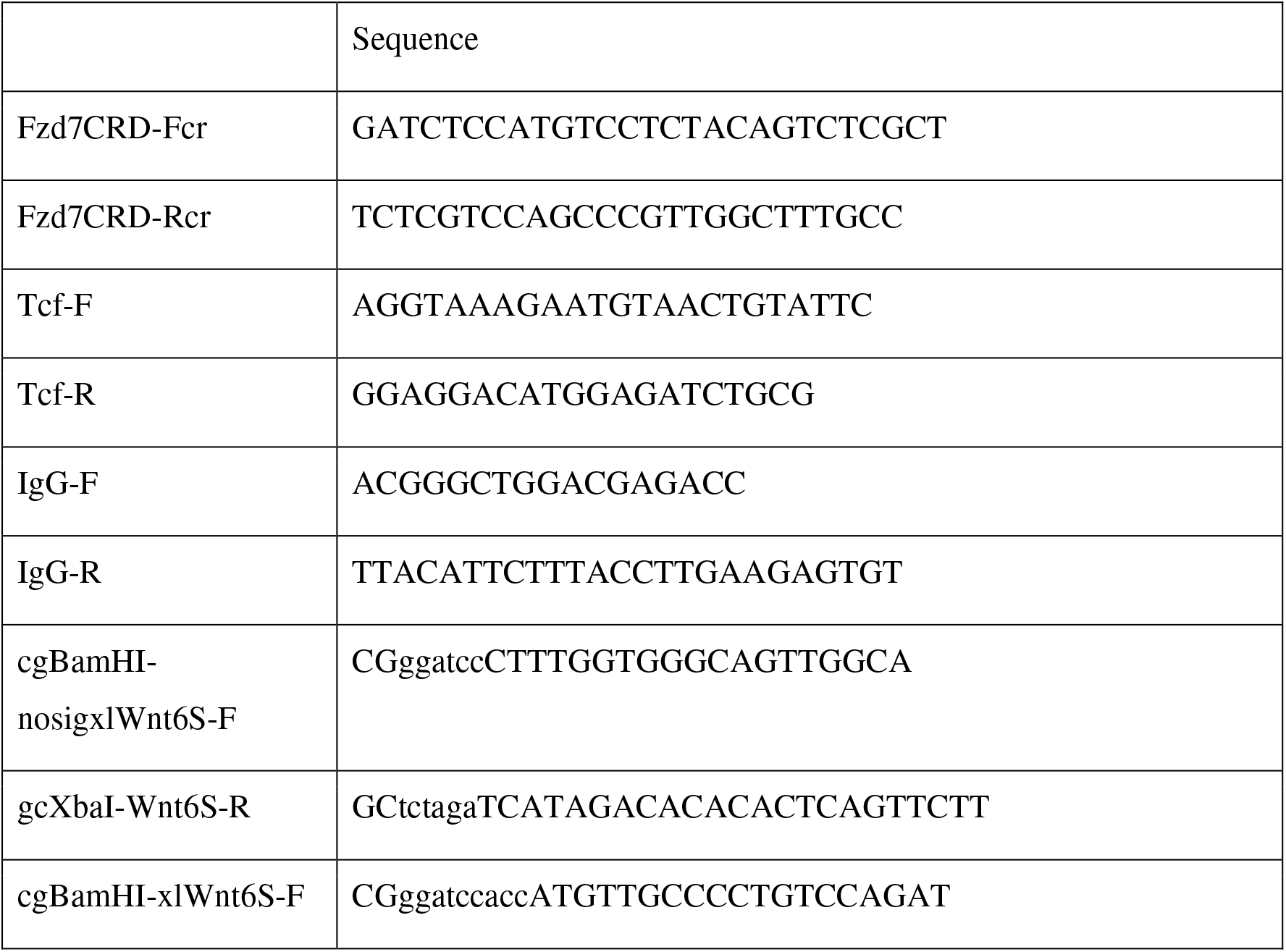
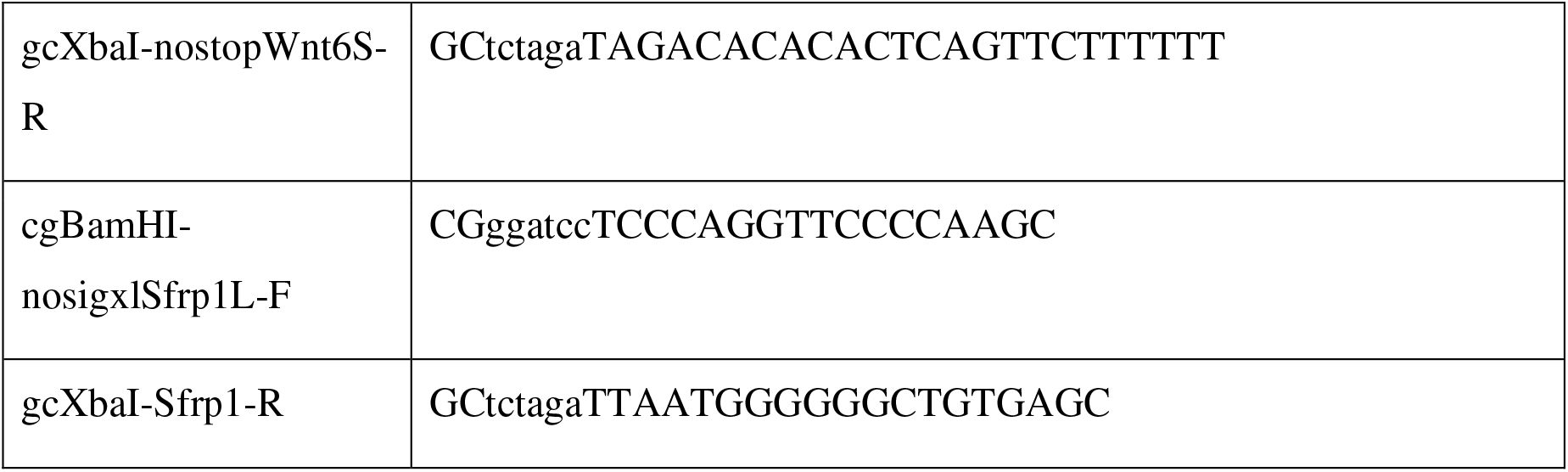
The primers that were used in this study. For construction cloning for mRNA synthesis:

For construction cloning of RNA in situ probes:

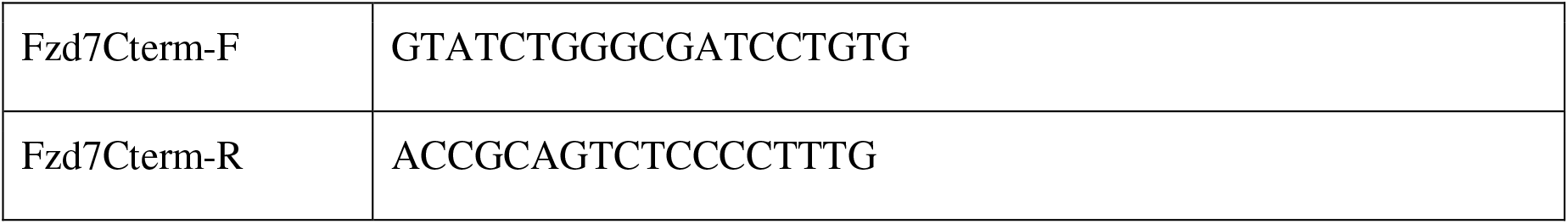

## Acknowledgements

Dr. Yukio Nakamura (Aberdeen University, UK) and Dr. Masanori Taira (Chuo University, Japan) for their help in the starting of this work. Dr. Takehiko Nakamura (Seikagaku Corporation, Japan) for NAH46 antibody and hybridoma. Dr. Osamu Yoshie (Kindai University, Japan) for HepSS-1 hybridoma. Dr. Makoto Matsuyama (Shigei Medical Research Institute, Japan) for the generation of NAH46 and HepSS-1 antibody from the hybridomas. This international collaboration was supported in part by Daiwa Anglo-Japanese Foundation (12969/13787 to T.Y.); with additional research support in Japan from MEXT/JSPS KAKENHI (19K16138 to T.Y., 18K06244/21K06183 to T.Y. and T.M.); and in the United Kingdom from BHF (RG/18/8/33673 to S.H.) and BBSRC (BB/N021924/1; BB/M001695/1 to S.H.). S.H. was a Royal Society/Leverhulme Trust Senior Research Fellow (SRF\R1\191017).

## Author contributions

T.Y., Y.K., T.M. and S.H. conceived this project; T.Y., Y.K. and B.A. performed experiments; T.Y., Y.O., and C.G. performed mathematical analysis. T.Y., Y.O., and S.H. wrote the manuscript with help from co-authors.

---

**Supplemental Fig. 1.**
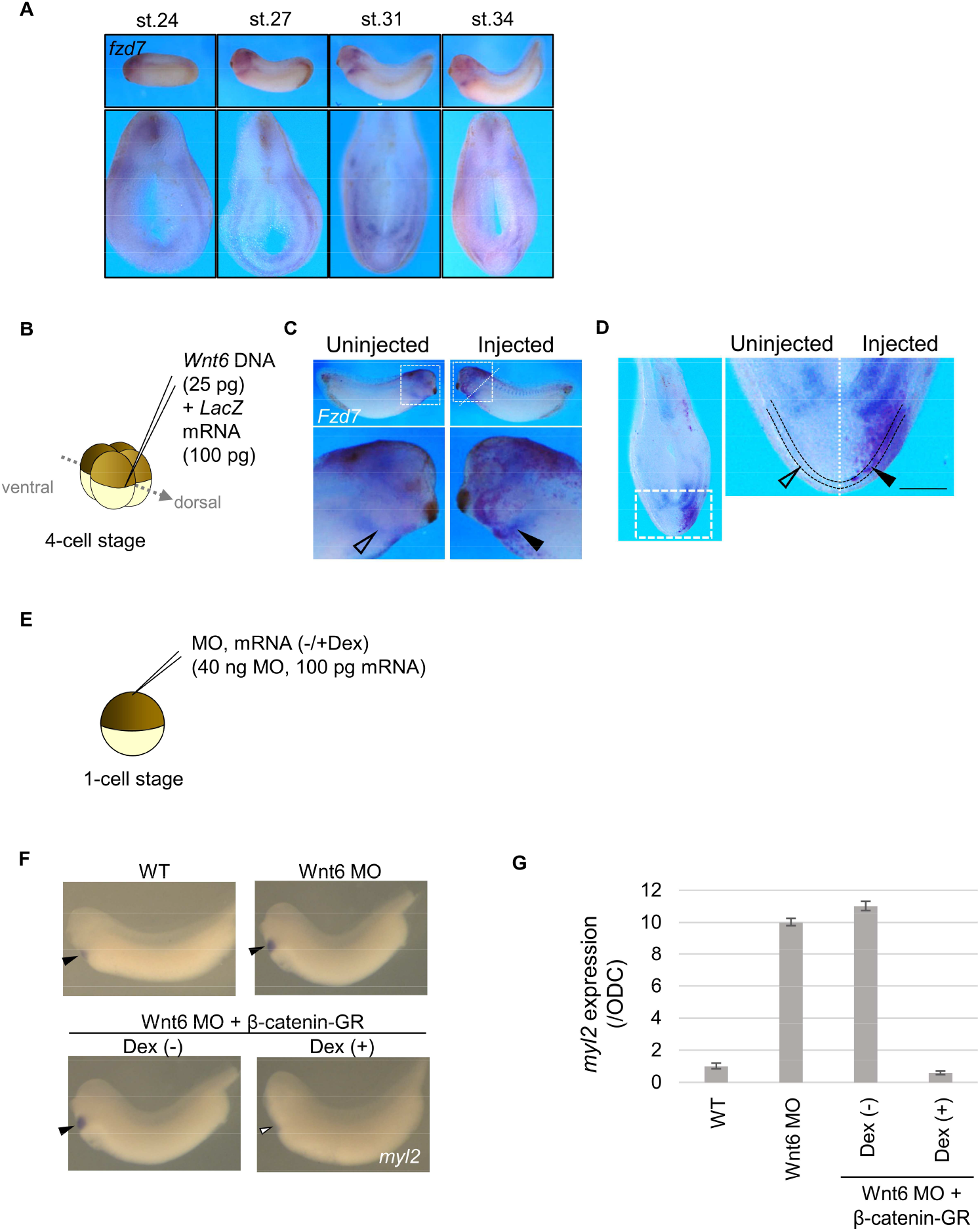
Wnt-dependent differentiation of the pericardium: Fzd7 expression. (A) *fzd7* expression during embryogenesis in *Xenopus*. The expression was broad at st. 24-27 but narrowed after st. 31. (B-D) Wnt6 DNA (25 pg) (see Materials & Methods) was injected into the marginal zone of one of the dorsal blastomeres at 4-cell stage as indicated (B). *lacZ* mRNA was coinjected as a tracer. In situ hybridization using *fzd7* probe (C-D). The specimen was cut at the position of dashed line to make hemi-section (C). *fzd7* expression area expanded in the injected side (arrowhead), compared with the uninjected side (open arrowhead) (D; n = 16/20). The pericardial region is the area between the dotted lines. Scale bar, 0.1 mm. (E-G) Wnt6 MO and mRNA of β-catenin-GR was injected into the animal pole at one-cell stage. β-catenin-GR was activated at the tailbud stage with Dex (E). The myocardium region (*myl2*-expressed region) was expanded by Wnt inhibition (n = 16/18) as reported and it was rescued by β-catenin activation (n = 19/19) (F). These results were confirmed by qRT-PCR (G; normalized by an internal control, ODC (ornithine decarboxylase), Bars represent SD.).

**Supplemental Fig. 2.**
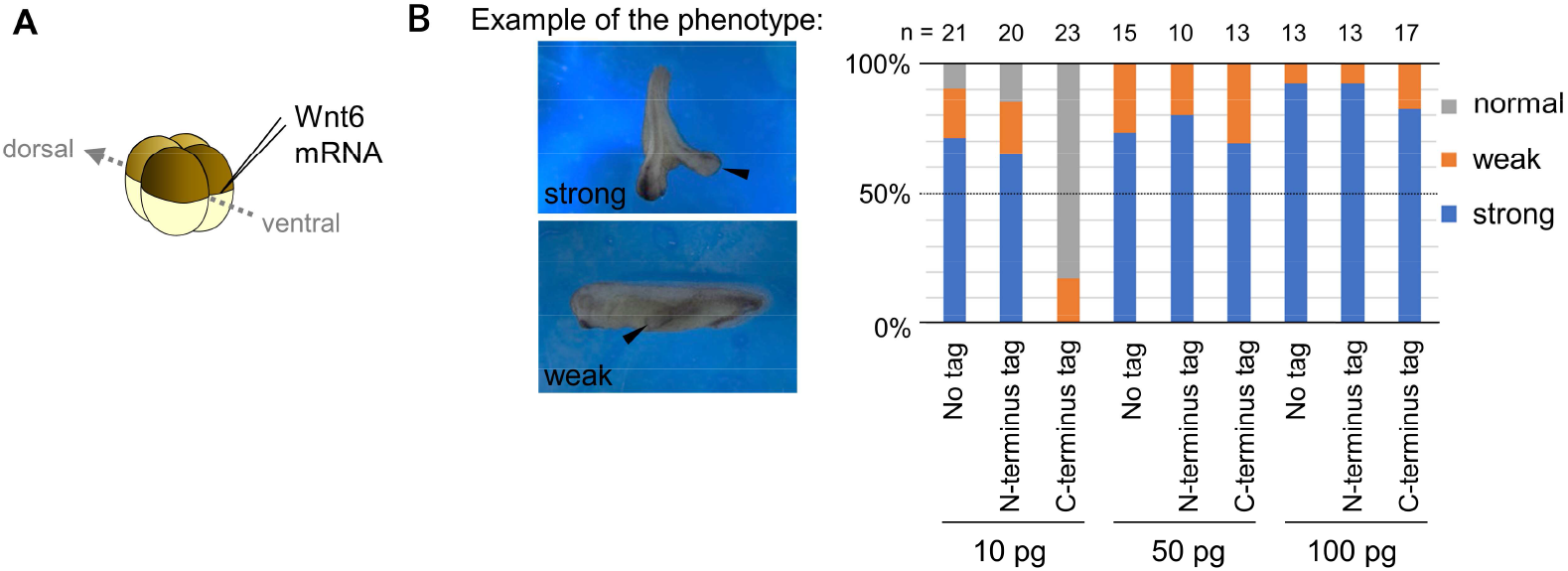
Biological activity of mVenus-tagged Wnt6 on secondary axis formation. (A) Each mRNA was injected into the marginal region of a ventral blastomere at 4-cell stage and the specimens were counted at the tailbud stage. (B) The biological activity of mVenus-tagged Wnt6 in different doses (the doses and the number of the specimen were indicated in the graph). The criteria of “strong” or “weak” is indicated in the left.

**Supplemental Fig. 3.**
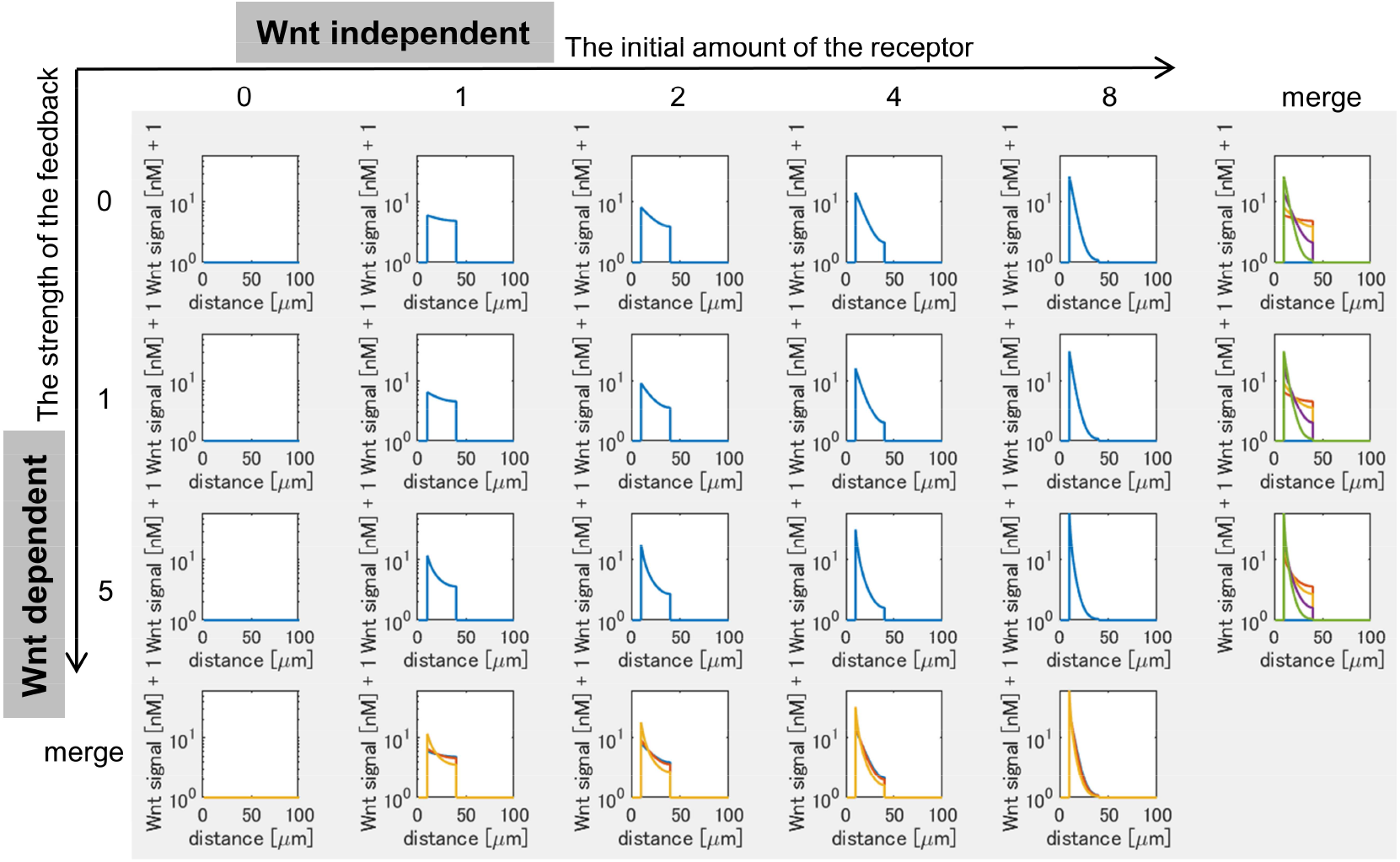
Mathematical modelling of Wnt activity in the heart region. Wnt activity was shown at t_max_ = 10^5^ sec (∼ 1 day after the onset of the simulation). The initial value of the receptor (the amount is Wnt-independent) increases toward the right, and the feedback strength (the amount is Wnt-dependent) increases toward the bottom. The numbers at the top and left of the figure indicate these relative amounts. In the merged figure, each line was displayed with different colors.

**Supplemental Fig. 4.**
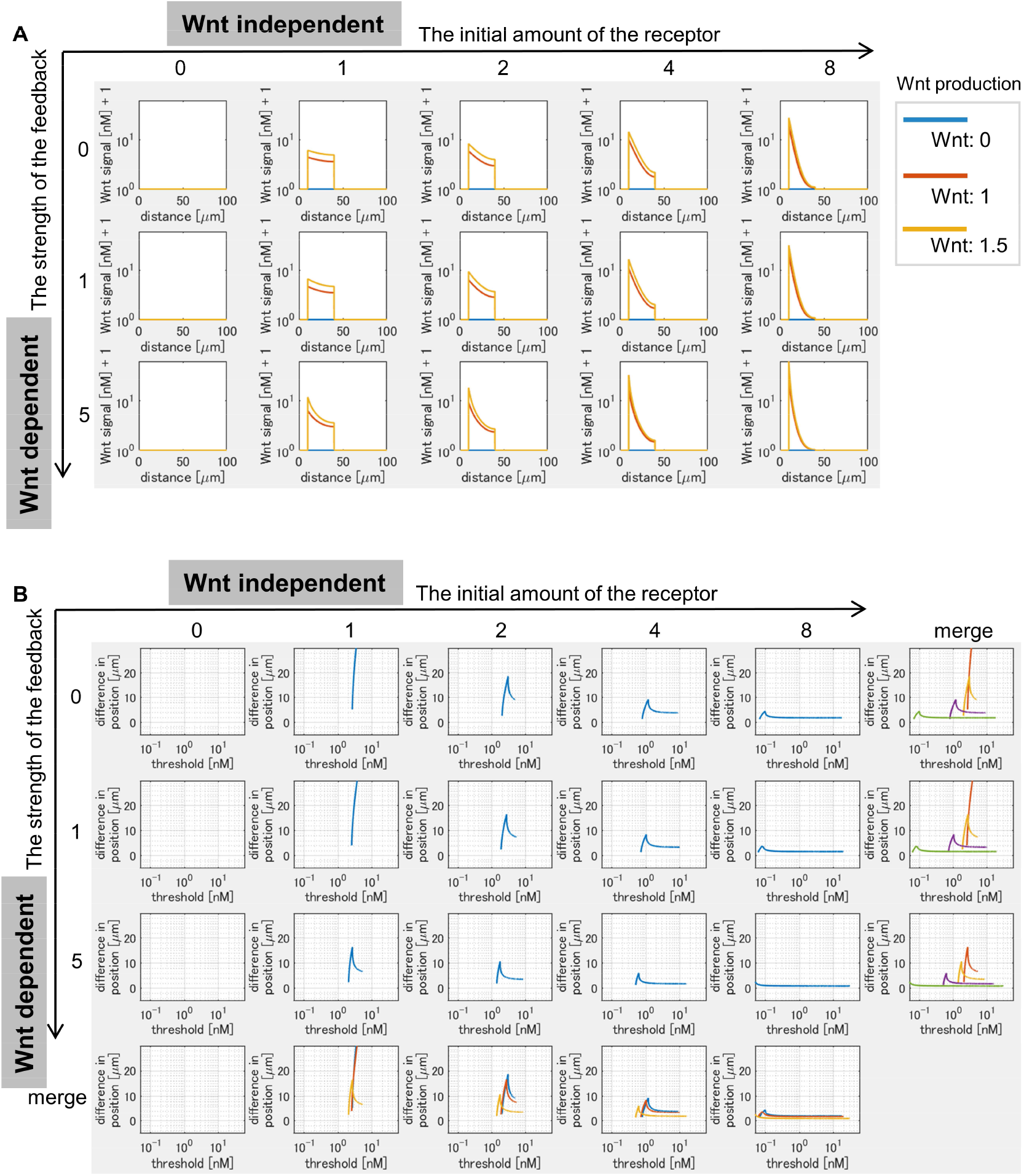
Changes in Wnt activity by fluctuating Wnt production. (A) Three lines in each graph shows Wnt activity with a different Wnt production rate at t_max_ = 10^5^ sec (∼ 1 day after the onset of the simulation). (B) The vertical axis shows the difference of boundary position when Wnt production rate is changed (1.0 to 1.5: 50% fluctuation in the ligand production). The horizontal axis indicates threshold values.

**Supplemental Fig. 5.**
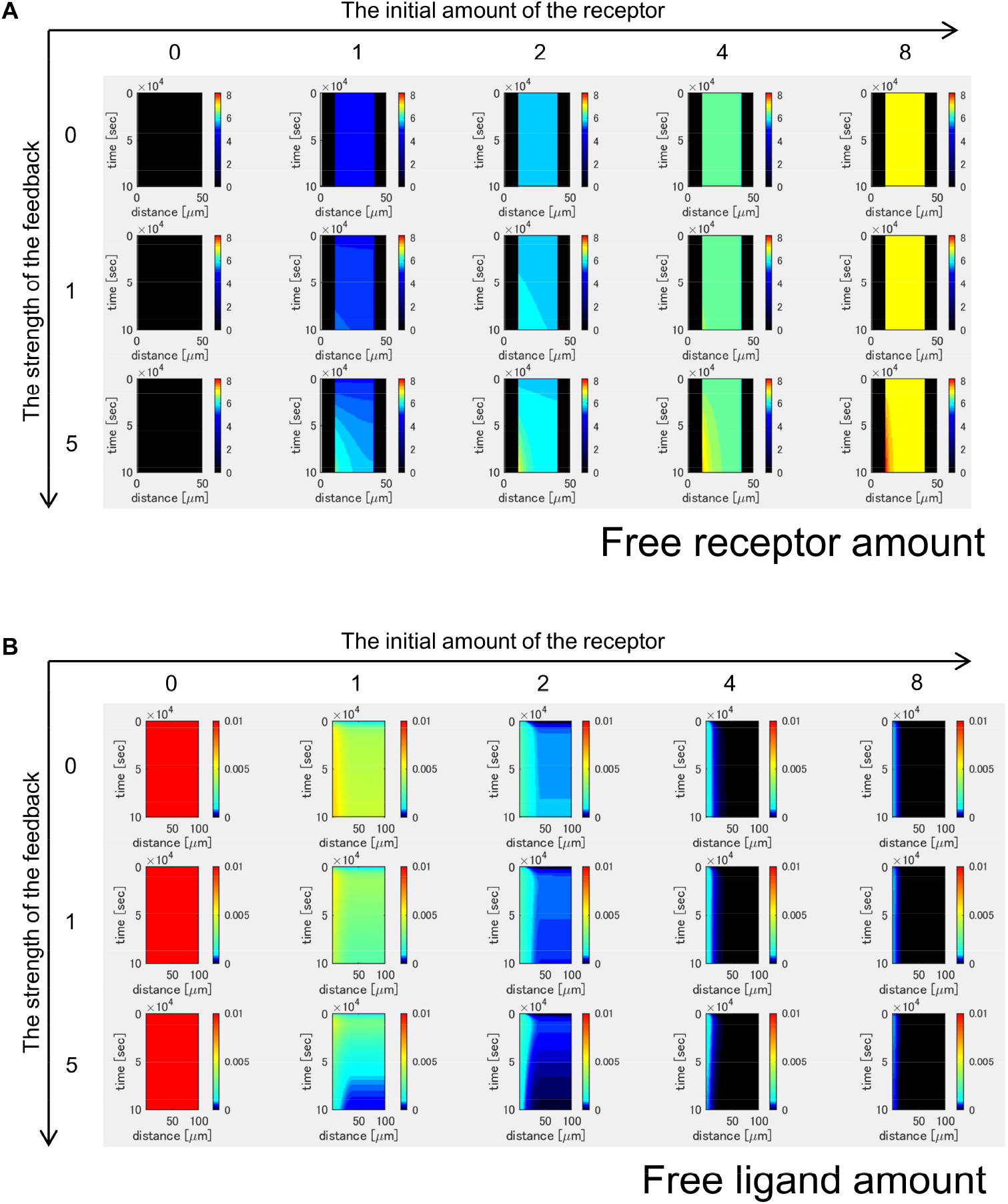
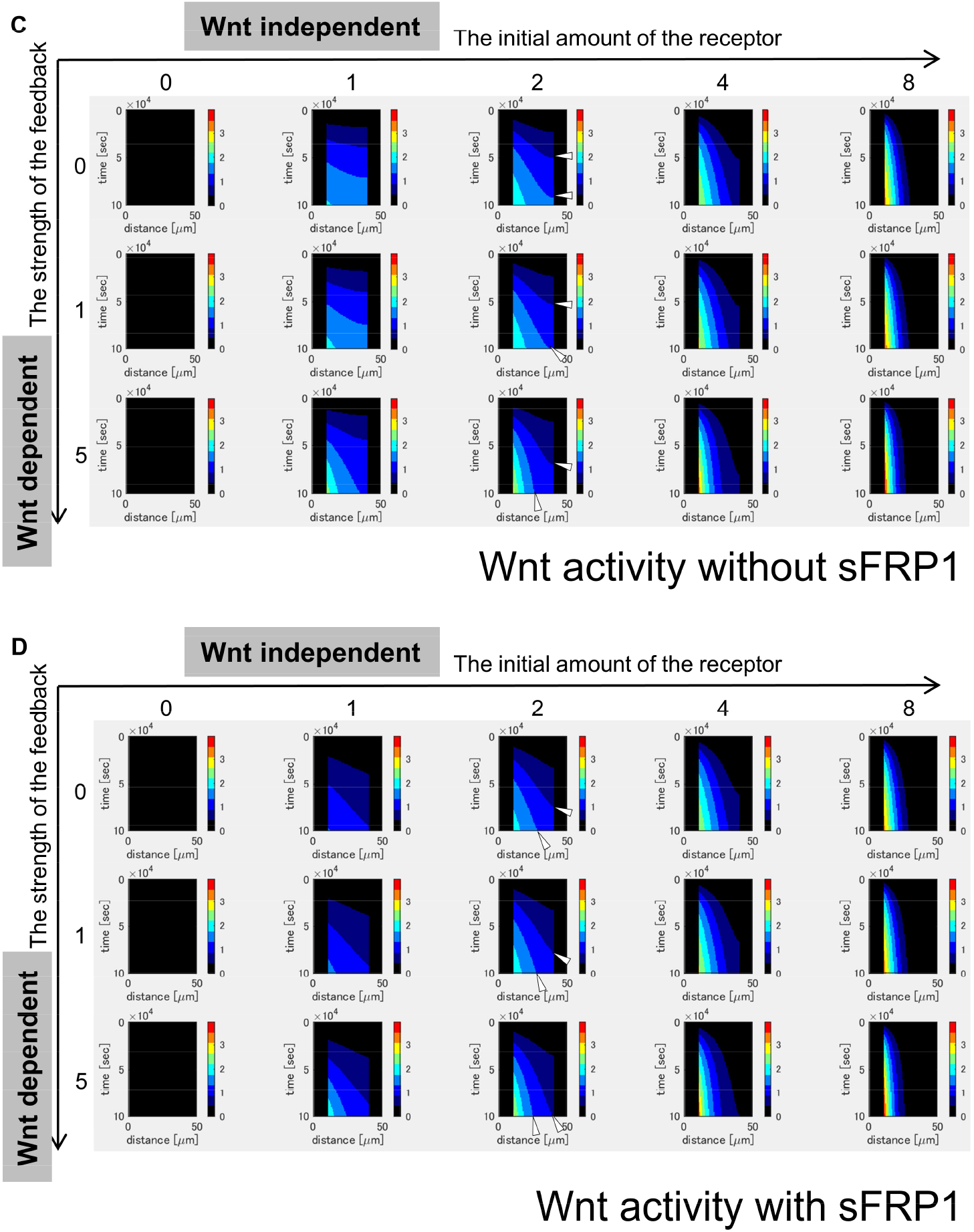
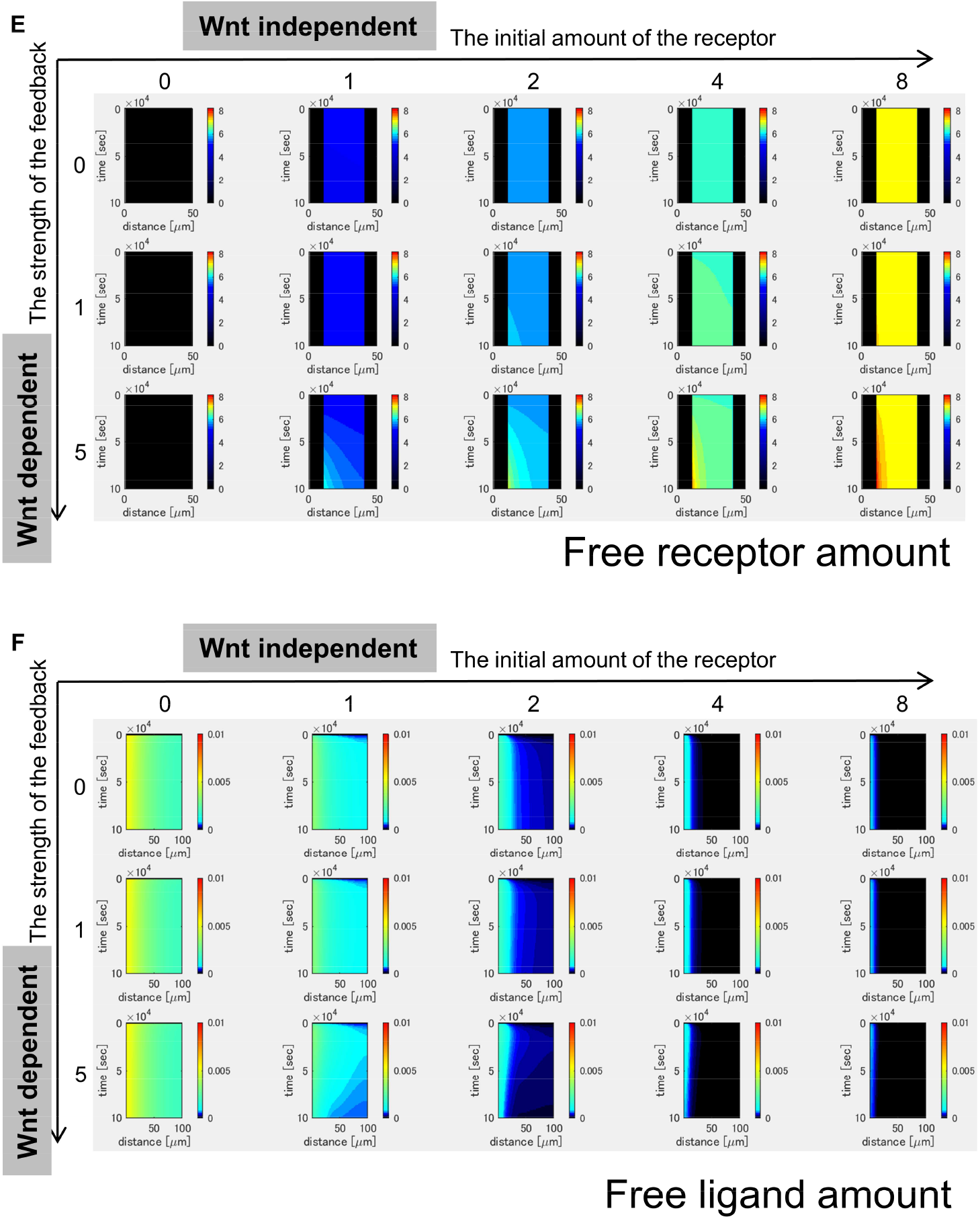
Heatmap image of Wnt signaling components and activity. The time change was shown as kymographs (Wnt production rate: 1.5 in Fig. 3B, Supplemental Fig. 4A), in which time proceeded from the upper to the bottom. (A) The time change of free receptor amount. The gradient can be observed with higher strength of the receptor feedback (for instance, feedback: 5, initial amount: 1). (B) The time change of free ligand amount. The amount was higher in low initial amount of the receptor and no feedback. With the feedback, however, the amount was gradually reduced (for instance, feedback: 5, initial amount: 1). (C-D) The time change of Wnt-signal activity without sFRP1(C) or with sFRP1 (D). Arrows indicates the axis of two colors boundary as an example. (E-F) The time change of free receptor amount (E) and free ligand amount (F) with sFRP1 condition.

**Supplemental Fig. 6.**
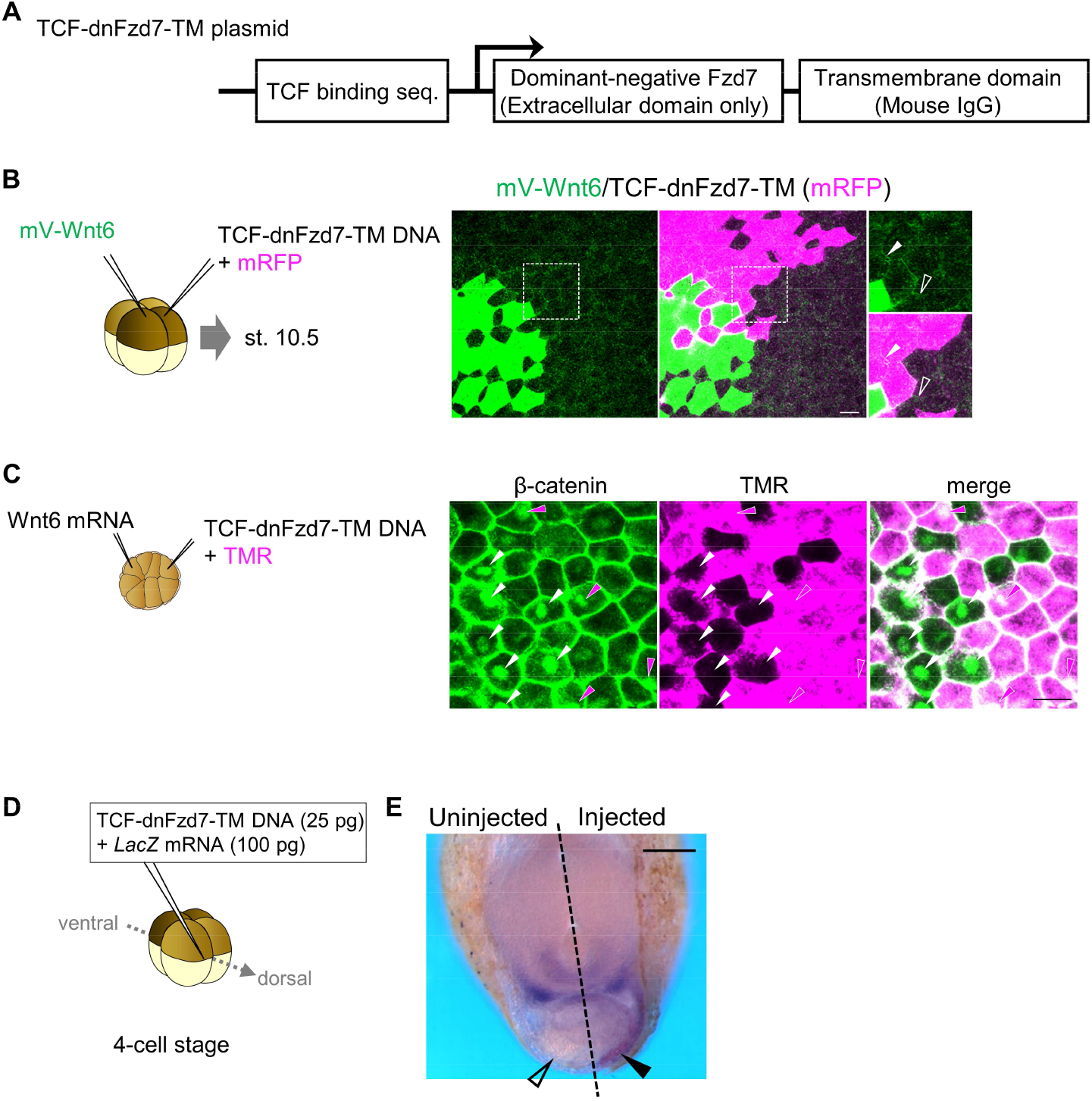
Receptor reduction expanded Wnt-signal-activated region. (A) The detail of the construct (TCF-dnFzd7-TM) inhibiting Fzd7 function in a Wnt-dependent manner. Luciferase sequence in TOPFlash plasmid was replaced with the sequence of dnFzd7, which is the extracellular domain of *fzd7*, and mouse IgG transmembrane. (B) mV-Wnt6 mRNA and TCF-dnFzd7-TM plasmid (with mRFP mRNA) were injected into different blastomeres at 4-cell stage (as indicated in the left). mV-Wnt6 signal was high (arrowhead) on the cell membrane between cells with TCF-dnFzd7-TM plasmid injected (magenta), compared to that between the intact cells (open arrowhead). This indicates dnFzd7-TM can accumulates Wnt6 in a cell autonomous manner. (C) Wnt6 mRNA and TCF-dnFzd7-TM plasmid (with a tracer, tetramethylrhodamine-dextran) were injected into different blastomeres at 16-cell stage (as indicated in the left). The hallmark of activated Wnt signaling, nuclear localization of β-catenin was reduced in the TCF-dnFzd7-TM injected cells (magenta region, compared to the intact cells (non-magenta region). Nuclear localizations of β-catenin were indicated by arrowheads (intact cells, white; dnFzd7-expressed cells, magenta). It should be noted that DNA injection generally induces mosaic expression, thus some of the injected cells express dnFzd7-TM. (D) TCF-dnFzd7-TM plasmid was injected into marginal region of dorsal blastomere at 4-cell stage. (E) In situ hybridization using Fzd7 intracellular domain probe. The probe recognizes the intracellular domain of *fzd7*, which the dominant negative *fzd7* lacks. Thus, it visualizes endogenous expression of full-length *fzd7* expression area (pericardium region), or Wnt-signal-activated region. Normal Fzd7 expression area, or Wnt-signal-activated region, is expanded (arrowhead) by injection of TCF-dnFzd7-TM plasmid (n = 11/11).

**Supplemental Fig. 7.**
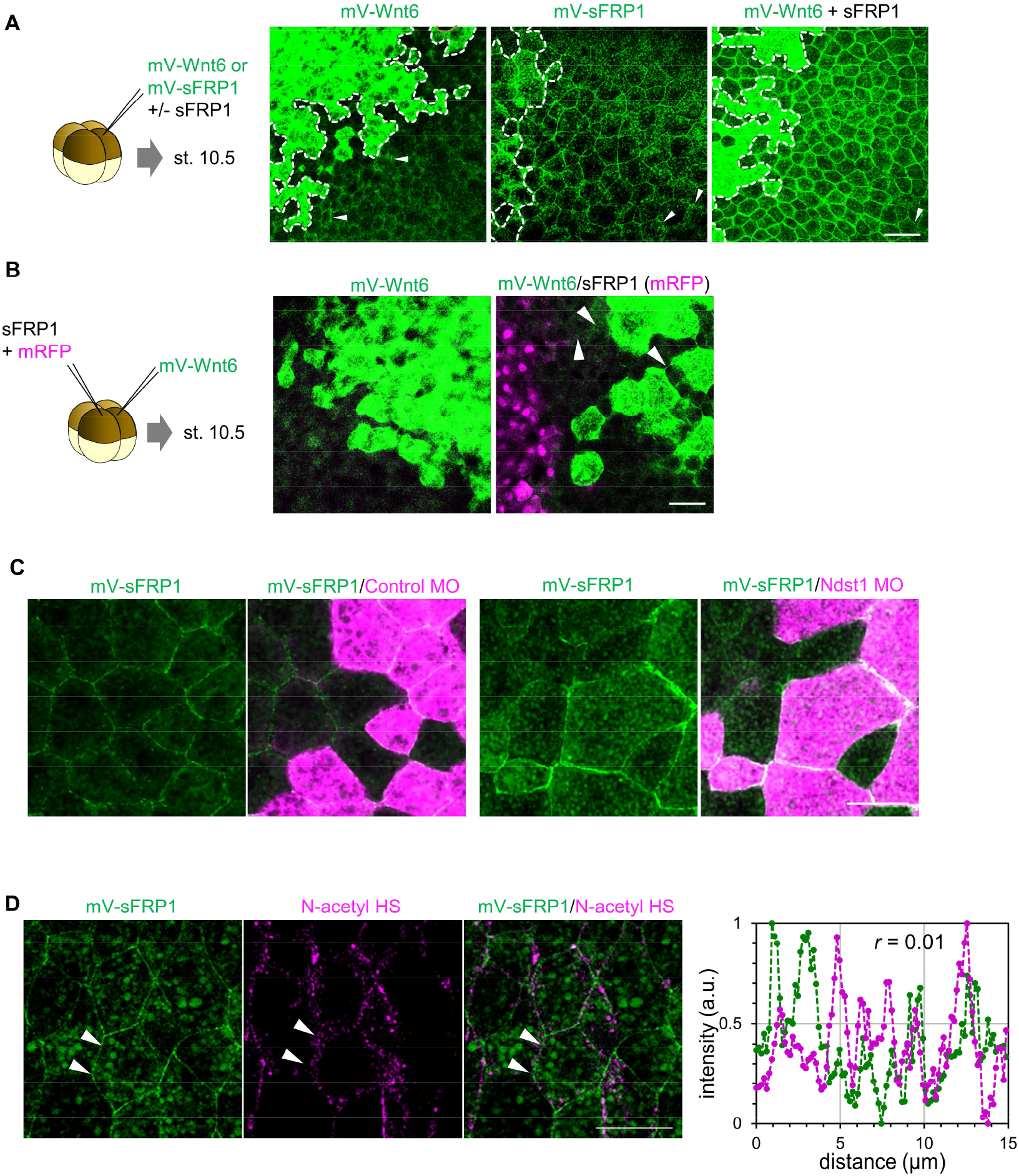
The distribution range and the role of sFRP1. (A) mV-Wnt6 with/without sFRP1 or mV-sFRP1 was injected into the ventral blastomere at 4-cell stage and observed at st. 10.5. The cells surrounded by dotted line indicates the source cells. Wnt6 distribution range was seemed to be short, whereas sFRP1 was long. sFRP1 can lengthen the distribution range of Wnt6. (B) The distribution of Wnt6 on the cell membrane was enhanced by sFRP1, which was expressed even at a distance from the source of Wnt6. mV-Wnt6 and sFRP1 were injected into different blastomeres. When mV-Wnt6 was injected, the distribution on the cell membrane other than the source cell was not clearly observed. However, the distribution can be seen when sFRP1 was injected (arrowheads). (C) sFRP1 was highly accumulated on *ndst1*-knockdowned cells. mV-sFRP1 mRNA and *ndst1* MO (with a tracer, TMR) were injected different blastomeres at 4-cell stage. (D) sFRP1 was not localized on N-sulfo-rich HS. mV-sFRP1 mRNA was injected into one blastomere at 4-cell stage. N-sulfo-rich HS was visualized by IHC. Signal intensity was showed in the graph, which was measured between two arrowheads (green, mV-sFRP1; magenta, N-sulfo-rich HS). Correlation coefficient (*r*) was shown in the graph. Scale bar: 50 μm (A,B), 30 μm (C,D).

